# Do chromosome rearrangements fix by genetic drift or natural selection? A test in *Brenthis* butterflies

**DOI:** 10.1101/2023.06.16.545248

**Authors:** Alexander Mackintosh, Roger Vila, Simon H. Martin, Derek Setter, Konrad Lohse

**Author notes:** These authors contributed equally to this work (Derek Setter, Konrad Lohse).

## Abstract

Large-scale chromosome rearrangements, such as fissions and fusions, are a common feature of eukaryote evolution. They can have considerable influence on the evolution of populations, yet it remains unclear exactly how rearrangements become established and eventually fix. Rearrangements could fix by genetic drift if they are weakly deleterious or neutral, or they may instead be favoured by positive natural selection. Here we compare genome assemblies of three closely related *Brenthis* butterfly species and characterise a complex history of fission and fusion rearrangements. An inferred demographic history of these species suggests that rearrangements became fixed in populations with large long-term effective size (*N_e_*). However, we also find large runs of homozygosity within individual genomes and show that a model of population structure with smaller local *N_e_* can reconcile these observations. Using a recently developed analytic framework for characterising hard selective sweeps, we find that chromosome fusions are not enriched for evidence of past sweeps compared to other regions of the genome. Nonetheless, one chromosome fusion in the *B. daphne* genome is associated with a valley of diversity where genealogical branch lengths are distorted, consistent with a selective sweep. Our results suggest that drift is a stronger force in these populations than suggested by overall genetic diversity, but that the fixation of strongly underdominant rearrangements remains unlikely. Additionally, although chromosome fusions do not typically exhibit signatures of selective sweeps, a single example raises the possibility that natural selection may sometimes play a role in their fixation.

## Introduction

### How do chromosome rearrangements fix?

Eukaryotic genomes vary widely in chromosome number and structure, i.e. karyotype. While closely related species often have similar karyotypes, there are also examples of considerable variation in chromosome number within genera (Hipp *et al*. 2009; Lukhtanov 2015) and even species (John and Hewitt 1970; Searle 1991; Zima *et al*. 1996). This variation is typically generated through chromosome rearrangements, with chromosome fissions and fusions resulting in increases and decreases in chromosome number, respectively. These rearrangements have been shown to promote speciation as well as influence the rate and distribution of recombination events (Bidau *et al*. 2001; Davey *et al*. 2017; Yoshida *et al*. 2023; Näsvall *et al*. 2023; Mackintosh *et al*. 2023), but our understanding of their role in evolution is limited by the fact that we do not know how they rise to high frequency in the first place. Heterozygosity for fissions or fusions can cause improper segregation during meiosis (White 1973), and so it is often suggested that new rearrangements are weakly deleterious and establish through strong genetic drift (Wilson *et al*. 1975; Bush *et al*. 1977). An alternative view is that rearrangements become fixed because they are favoured by natural selection (Bickham and Baker 1979; Qumsiyeh and Handal 2022), but there is currently limited empirical evidence to support this.

There are a number of different ways for a fission or fusion to be advantageous. For example, a chromosome fusion can increase linkage disequilibrium (LD) between coadapted alleles, leading to enhanced local adaptation and fixation of the rearrangement (Fisher 1930; Charlesworth 1983; Guerrero and Kirkpatrick 2014). There are examples of fused chromosomes that are enriched for adaptive loci (Wellband *et al*. 2019; Liu *et al*. 2022), but it is unclear what fraction of these variants predate the rearrangements and potentially contributed to their fixation. Rearrangements can also have direct effects on gene expression, either through changes in genome positioning within the nucleus (Di Stefano *et al*. 2020) or if breakpoints occur within a gene body or regulatory element (Harewood and Fraser 2014). While most changes in gene expression are likely deleterious, any beneficial changes could lead to the spread of a rearrangement. Meiotic drive is another mechanism by which chromosome rearrangements could rapidly increase in frequency. This process involves drive alleles that are transmitted to gametes more than 50% of the time and typically occurs within asymmetric meiosis (Pardo-Manuel de Villena and Sapienza 2001b). Chromosome rearrangements with differences in centromere size or form can act as drive alleles which leads to their fixation (Pardo-Manuel de Villena and Sapienza 2001a; Stewart *et al*. 2019). Although this process is primarily associated with monocentric chromosomes (i.e. those with a single centromere) it has also been suggested to occur in organisms with holocentric chromosomes, such as nematodes and Lepidoptera, where centromeres are not localised (Burěs and Zedek 2014).

While the processes described above are certainly possible, the fixation of fissions and fusions may not be adaptive at all. Instead, a rearrangement could fix entirely through genetic drift (Wright 1941). This may be the case if meiosis is robust to the risk of unbalanced segregation associated with heterokaryotypes (Borodin *et al*. 2019). Even if a rearrangement does confer a fitness cost, strong drift and inbreeding in small populations could still lead to its fixation (Wright 1941; Lande 1979). Under this scenario, one would expect more rearrangements involving Y/W chromosomes than those involving X/Z chromosomes, due to the approximately three-fold difference in effective population size (*N_e_*). Pennell *et al*. (2015) tested this prediction and found that Y/W-autosome fusions are indeed significantly more common than X/Z-autosome fusions in fish and squamate reptiles, though not in mammals. They therefore suggest that sex-autosome fusions are often weakly deleterious and fix through genetic drift. While the same could be true for fissions and autosome-autosome fusions, it is unclear whether all of these rearrangements have similar fitness effects.

### Inferring selective sweeps

If fissions and fusions rise in frequency due to natural selection, sites that are tightly linked to recent rearrangements will show evidence of selective sweeps. This process, in which a beneficial allele increases rapidly in frequency and nearby alleles ’hitchhike’ with it, leaves a signature in population genomic data that can be used to infer past selection (Maynard Smith and Haigh 1974). A variety of methods have been developed for sweep inference, often making use of different types of genomic data, such as allele frequencies (Nielsen *et al*. 2005), patterns of haplotype similarity (Garud *et al*. 2015; Harris and DeGiorgio 2020), or even reconstructed ancestral recombination graphs (Stern *et al*. 2019; Hejase *et al*. 2022). One limitation shared by a number of methods is the assumption that the modelled selective sweep has completed very recently. This limits the power to detect and accurately characterise even strong sweeps that happened deeper in time.

Recently, Bisschop *et al*. (2021) showed that, for small sample sizes, the joint distribution of genealogical branch lengths can be derived under an approximate model of a selective sweep. This allows the calculation of composite likelihoods from mutation configurations in short sequence blocks, in particular the blockwise site frequency spectrum (bSFS; Bunnefeld *et al*. 2015). Importantly, this analytic framework can be used to infer and characterise sweeps that happened further back in time (i.e. *>* 0.1*N_e_* but *<* 4*N_e_* generations ago) by treating the sweep as a discrete event that is both preceded and followed by a neutral coalescent process (Bisschop *et al*. 2021). This inference method can therefore be used to test whether natural selection has acted on certain regions of the genome, even if the selective events are relatively old.

### Overview

Here we use the fast rate of chromosome evolution in *Brenthis* fritillary butterflies to investigate how chromosome fissions and fusions evolve. Previous work has shown that chromosome numbers vary substantially among the four species in this genus (Saitoh 1986; Saitoh and Lukhtanov 1988; Saitoh 1991; Pazhenkova and Lukhtanov 2019; Mackintosh *et al*. 2022) and that this variation is due to chromosome rearrangements (Mackintosh *et al*. 2023), rather than differences in ploidy or supernumerary chromosomes. First, we describe a newly generated chromosome-level genome assembly for *Brenthis hecate*. This species has a much larger number of chromosomes (*n_c_* = 34) than *B. daphne* (*n_c_* = 12-13) or *B. ino* (*n_c_* = 13-14), implying a history of rapid rearrangement. Secondly, we compare the genomes of these three *Brenthis* species with publicly available genome assemblies of two other fritillary butterfly species in the tribe Argynnini. Using a maximum parsimony method, we show that almost all rearrangements are confined to the genus *Brenthis*. Thirdly, we use whole genome resequence data for all three *Brenthis* species to estimate their demographic history. This allows inferred rearrangements to be placed within the context of species divergence times and effective population sizes. Finally, we investigate whether chromosome fusions, the most common rearrangement type in our dataset, have fixed by natural selection. For each of 12 potentially recent chromosome fusions, we use the analytical framework of Bisschop *et al*. (2021) to estimate support for a sweep model, as well as the time since the sweep and the strength of selection.

## Materials and methods

### Sampling and sequencing

Butterflies were collected by hand netting and frozen from live in a -80 freezer. We performed a high molecular weight (HMW) DNA extraction for one *Brenthis hecate* individual (ES BH 1412; Table S1) using a salting-out protocol (see Mackintosh *et al*. 2022 for details). For four other *B. hecate* individuals (Table S1), DNA was extracted from Ethanol preserved samples using a Qiagen DNeasy Blood & Tissue kit, following the manufacturer’s instructions. Edinburgh Genomics (EG) prepared TruSeq Nano gel free libraries from all five DNA extractions and sequenced them on an Illumina NovaSeq 6000. EG also generated a SMRTbell sequencing library from the HMW DNA and sequenced it on a Pacbio Sequel I instrument. A sixth individual (ES BH 1411; Table S1) was used for chromatin conformation capture (HiC) sequencing. EG performed the HiC reaction using an Arima-HiC kit, following the manufacturer’s instructions for flash frozen animal tissue, and generated a TruSeq library which was sequenced on an Illumina NovaSeq 6000.

### Genome assembly

We generated a reference genome for *B. hecate* by assembling Pacbio continuous long reads with Nextdenovo v2.4.0 (Hu *et al*. 2023). The contig sequences were polished with Illumina short-reads from the same individual using Hapo-G v1.1 (Aury and Istace 2021). We identified and removed haplotypic duplicates and contigs deriving from other organisms using purge dups v1.2.5 (Guan *et al*. 2020) and Blobtools v1.1.1 (Laetsch and Blaxter 2017), respectively. We mapped HiC data to the contigs with bwa-mem v0.7.17 (Li 2013) and then used YaHS v1.1a.2 and juicebox v1.11.08 to scaffold the assembly into chromosome-level sequences (Zhou *et al*. 2023; Robinson *et al*. 2018).

### Synteny analysis

We compared synteny between five genome assemblies of butterfly species in the tribe Argynini: *Brenthis hecate*, *Brenthis ino* (Mackintosh *et al*. 2022), *Brenthis daphne* (Mackintosh *et al*. 2023), *Fabriciana adippe* (Lohse *et al*. 2022b), and *Boloria selene* (Lohse *et al*. 2022a). Pairwise alignment of assemblies were performed with minimap2 v2.17 (Li 2018) and differences in synteny were visualised by plotting high quality alignments (mapping quality of 60 and length *>*= 50 kb). We found that the genome sequence of *B. selene* has low sequence identity to the other genomes, resulting in few nucleotide alignments. We therefore identified BUSCO genes in all five assemblies (lepidoptera odb20, BUSCO v5.3.2, Simão *et al*. 2015) and used the location of these BUSCO genes to visualise synteny between the *B. selene* genome and the others.

We estimated the number of fission and fusion rearrangements across the phylogeny of these species using syngraph (https://github.com/DRL/syngraph). We included an additional nymphalid genome assembly in this analysis (*Nymphalis polychloros*, Lohse *et al*. 2021) as an outgroup. BUSCO genes were used as markers and the minimum number of markers for a rearrangement to be reported was set to five. We used the tabulated output of syngraph, as well as the paf files generated by minimap2, to identity approximate positions of chromosome fusion points.

### Demographic inference

To infer a demographic history for the three *Brenthis* species, we mapped whole genome resequencing (WGS) data to the *F. adippe* reference genome. This included data for five *B. hecate* individuals (Table S1), as well as seven *B. daphne* and six *B. ino* individuals that were originally analysed in Mackintosh *et al*. (2023). Individuals were sampled from across the Palearctic (Table S1, see Figure 1 in Mackintosh *et al*. 2023), including different glacial refugia. WGS data were trimmed with fastp v0.2.1 (Chen *et al*. 2018) and mapped with bwa-mem. Variants were called with freebayes v1.3.2 (Garrison and Marth 2012) and filtered with gIMble preprocess (Laetsch *et al*. 2022) using the following options: `--snpgap 2 --min qual 10 --min depth 8 --max depth 5`. We applied an additional filter to remove SNPs where *>* 70% of individuals were heterozygous, as these are likely due to alignment of paralogous sequence. We annotated genes in the *F. adippe* genome (see Supplementary Methods) and used this to restrict our analysis to four-fold degenerate (4D) sites. Given 295,730 SNPs, as well as a total count of 4D sites callable across all individuals (2,487,949), we generated an unfolded three dimensional site frequency spectrum (3D-SFS) using get 3D SFS.py (see Data accessibility). The ancestral state at each SNP was assigned using the reference (*F. adippe*) allele. Demographic modelling was performed with fastsimcoal2 (fsc27093) (Excoffier *et al*. 2021). We first fit a maximally complex model of divergence and gene flow between the three *Brenthis* species which included two split times, six effective population sizes, and eight asymmetrical migration rates (16 parameters total). Given the results across replicate analyses, we identified migration rates that could be made symmetrical as well as those that could be set to zero. We then considered three competing models, each with a total of 12 parameters, and identified the one with the greatest composite likelihood. Demographic parameter estimates were scaled using a *de novo* mutation rate of 2.9 *×* 10*^−^*^9^ (Keightley *et al*. 2015). A detailed description of the maximum likelihood model selection with fastsimcoal2 can be found in the Supplementary Methods.

**Figure 1:**
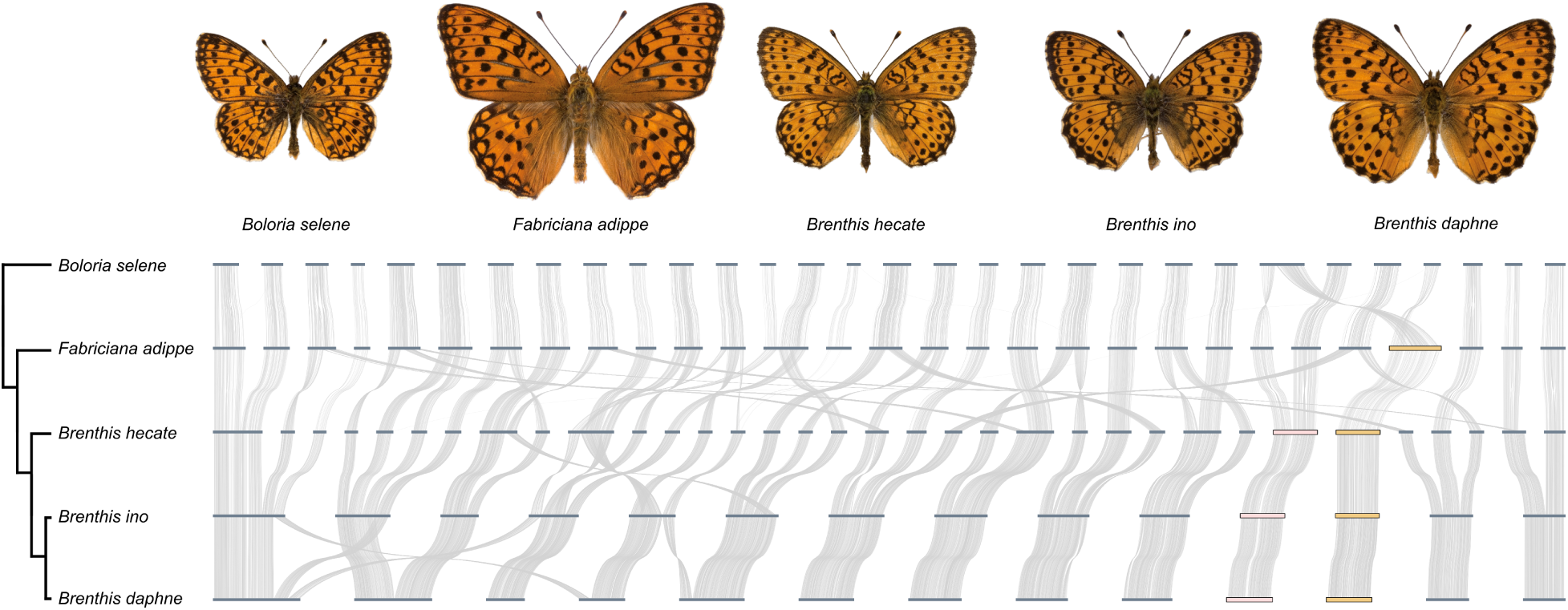
Synteny relationships between genomes of three *Brenthis* species, as well as species from two related genera. Top: Uppersides of male butterflies representing each of the five species. Bottom: A tree showing phylogenetic relationships between the species. The topology is from Chazot *et al*. (2021) and the plotted branch lengths are not to scale. Whole genome alignments are shown to the right of the tree. Thick horizontal bars are chromosomes and curved lines are nucleotide alignments, or, in the case of *B. selene* and *F. adippe*, shared BUSCO genes. Two sets of orthologous chromosomes are highlighted towards the right of the plot: an autosomal chromosome shared by all three *Brenthis* species (pink) and the Z chromosome that is shared by all *Brenthis* species and *F. adippe* (orange).

### Identifying runs of homozygosity

We identified runs of homozygosity (ROH) in each *Brenthis* individual to gain more information about genetic drift in the recent past of these species. To do this, we mapped WGS data for each *Brenthis* species to the corresponding (species-specific) reference genome with bwa-mem. Variants were called within each species using freebayes and filtered with gIMble preprocess: `--snpgap 2 --min qual 10 --min depth 8 --max depth 1.5`. We restricted SNPs in the VCF to non-repeat regions where all individuals had a genotype. We then identified runs of homozygosity (ROH) in each individual using plink v1.90b6.18 (Purcell *et al*. 2007) with the following options: `--homozyg-window-snp 1000 --homozyg-window-het 10 --homozyg-window-threshold 0.001 --homozyg-kb 100`.

### Fitting models of intraspecific population structure

For each *Brenthis* species we estimated demographic parameters under a finite-island model (Maruyama 1970). We fit the model to three summary statistics: per-site heterozygosity (*H*), pairwise intraspecific divergence (*d*_xy_) between individuals sampled from different locations in Europe, and the proportion of 1 Mb windows covered by a ROH (which we call *W*_roh_). For a given species, we averaged these statistics across all individuals / pairwise comparisons. We used the expected time of coalescence for lineages sampled within and between demes (Nagylaki 1982; Strobeck 1987; Wakeley 1999) to calculate the expected *H* and *d*_xy_ respectively. We estimated the probability of observing a 1 Mb window covered by a ROH as the probability that, for two lineages sampled from the same deme, the first event backwards in time is coalescence rather than migration or recombination within the window. These calculations assume equal recombination and mutation rates (*µ* = *r* = 2.9 *×* 10*^−^*^9^). We then inferred the parameters of the finite-island model (*N_e_*, the number of demes and the migration rate *m*) by maximising the fit to the observed *H*, *d*_xy_ and *W*_roh_. Model fitting was performed in a *Mathematica* notebook (finite island model.nb, see Data accessibility).

### Inferring selective sweeps

We used the same species-specific filtered VCF files described above as data for inferring selective sweeps around chromosome fusions. We annotated genes in the assemblies (see Supplementary Methods) and removed SNPs within exons (+/- 10 bases), i.e. we only consider variation within intronic or intergenic sequence. We chose to analyse *n* = 4 diploids for each species, selecting the set of individuals that minimised pairwise intraspecific *F_st_*. We analysed 12 chromosome fusions, that each are private to one of the *Brenthis* species, i.e. we did not consider fusions shared by multiple species which likely fixed many generations ago. Two of the 12 fusions were not inferred by the maximum parsimony method described above. Syngraph inferred a single ancient fusion and then a subsequent fission in *B. daphne*. Independent fusions in *B. hecate* and *B. ino* are equally parsimonious and supported by the fact that different chromosome ends are involved in each case. We therefore include these potential fusions in our analysis. We summarised the sequence variation surrounding each fusion in terms of the blockwise site frequency spectrum (bSFS; Bunnefeld *et al*. 2015), setting a block size so that the average block contained 1.5 segregating sites. We used six lineage bSFS.py (see Data accessibility) to record the folded bSFS for six lineages by considering all possible sets of three diploids from *n* = 4. We then applied a *k_max_* value of 2 using format blocks.py (see Data accessibility), i.e. we recorded exact mutation counts up to a value of 2 in each block and any count greater was summarised as *>* 2. We implemented the sweep inference method of Bisschop *et al*. (2021) in *Mathematica* (see Data accessibility). For a given point in the genome, we estimated the composite likelihood of a neutral model and a selective sweep model given the bSFS counts in the surrounding megabase. We normalised the difference in composite likelihood (Δ ln *CL*) by the number of blocks to allow comparisons between 1 Mb regions with a different number of blocks. In cases where chromosome fusion points could only be narrowed down to intervals spanning *>* 5 kb, we sampled points every 5 kb and reported parameter values for the point with the greatest Δ ln *CL* (Table 1). As a comparison, we also fit sweep models to points sampled from a non-rearranged chromosome (Figure 1). Additional details of the model fitting procedure can be found in the Supplementary Methods.

**Table 1:** Maximum composite likelihood parameter estimates for selective sweeps around chromosome fusions in *Brenthis* butterflies. The sweep with the greatest statistical support is highlighted in bold.

| **Taxon** | **Chr** | **Position**<br>**(Mb)** | $Log_{10}(\alpha)$ | $T_a$ | $\Delta \ln CL$ **per-**<br>**block** |
| --- | --- | --- | --- | --- | --- |
| *B. daphne* | 1 | 36.7 | -2.29 | 1.00 | .000 |
| *B. daphne* | 1 | 45.2 | -4.71 | 0.10 | .013 |
| ***B. daphne*** | **2** | **23.9** | **-5.67** | **0.08** | **.123** |
| *B. daphne* | 3 | 7.4 | -5.70 | 0.09 | .086 |
| *B. daphne* | 8 | 6.7 | -5.70 | 0.25 | .016 |
| *B. hecate* | 2 | 6.8 | -4.60 | 0.35 | .007 |
| *B. hecate* | 2 | 19.4 | -5.70 | 0.89 | .012 |
| *B. hecate* | 5 | 15.0 | -4.08 | 1.00 | .000 |
| *B. ino* | 1 | 6.6 | -4.43 | 1.00 | .000 |
| *B. ino* | 3 | 24.1 | -5.70 | 1.00 | .008 |
| *B. ino* | 8 | 22.2 | -5.70 | 1.00 | .004 |
| *B. ino* | 9 | 7.4 | -5.03 | 1.00 | .000 |

### Simulations

To quantify the power and accuracy of sweep inference based on the bSFS, we performed coalescent simulations with msprime v1.0.2 (Baumdicker *et al*. 2021) and applied the sweep inference scheme to the simulated data. Three different scenarios were simulated: a strong selective sweep (*s* = 0.005, *T_a_*= 250,000 generations ago, with *N_e_*= 500,000), neutral evolution (*N_e_*= 500,000), and neutral evolution in a population with a similar demographic history to *B. daphne* (as inferred by fastsimcoal2). The mutation and recombination rates were set to *µ* = *r* = 2.9 *×* 10*^−^*^9^ per-site per-generation. Each simulation was replicated 100 times, where a single replicate consisted of a 1 Mb sequence sampled for *n* = 4 diploids.

### Statistical analysis

We used resampling tests to evaluate whether chromosome fusions are enriched for selective sweeps when compared to loci elsewhere in the genome. We measured two statistics – the number of fusions with putative sweeps and the sum of Δ ln *CL* across all fusions in each species – and compared these with points sampled from a non-rearranged chromosome (Figure 1). Resampling was species-specific, i.e. we sampled the same number of points as fusions analysed for each species. We generated 100,000 random sample sets and calculated one-tailed p-values as the proportion of samples with values greater than our observed statistics.

## Results

### A genome assembly of *Brenthis hecate*

We generated a chromosome-level genome assembly for *Brenthis hecate* using a combination of Pacbio long-reads, Illumina short-reads, and HiC data (Figure S1). The assembly is 408.76 Mb in length, with a scaffold N50 of 12.79 Mb and a contig N50 of 5.86 Mb. Of the 45 sequences in the assembly, 34 are chromosome-level (herafter simply referred to as chromosomes), whereas the remaining 11 are contigs that could not be scaffolded (15 - 104 kb in length, totalling 409 kb). The chromosomes show a bimodal distribution in size (Figure S1), with seven large chromosomes (21.52 - 29.02 Mb) and 27 smaller chromosomes (6.55 - 14.04 Mb). The number of chromosomes in the *B. hecate* genome assembly (*n_c_* = 34) is consistent with reports of spermatocytes sampled from both France and Siberia (de Lesse 1961; Saitoh and Lukhtanov 1988). The genome sizes of *B. hecate*, *B. daphne* and *B. ino* are all similar: 409, 419, and 412 Mb, respectively.

### Synteny between Argynnini butterfly species

To characterise chromosome rearrangements, we performed whole genome alignments between the three *Brenthis* species, as well as genome assemblies from two other fritillary butterfly genera in the tribe Argynnini. The whole genome alignments show that the two outgroup species, *Fabriciana adippe* and *Boloria selene*, have very similar genome/chromosome organisation (Figure 1). By contrast, genomes of the *Brenthis* species show evidence for many rearrangements (Figure 1).

We next placed fission and fusions events on the phylogeny (*Brenthis sp.* and outgroups) using a maximum parsimony method (see Methods). Of the 53 inferred rearrangements, 50 are found on branches leading to *Brenthis* species or their most recent common ancestors. The branch with the greatest number of inferred rearrangements (11 fissions and 9 fusions) is that leading to the common ancestor of the three *Brenthis* species. Closer to the present, 14 fusion rearrangements are estimated on the branch ancestral to *B. daphne* and *B. ino*, while five fissions and two fusions are estimated on the branch leading to *B. hecate*. These rearrangements explain the large difference in chromosome number between these species. We also infer one fission and five fusions on the branch leading to *B. daphne* and three fusions on the branch leading to *B. ino*. Together these rearrangements form a complex history of genome ’reshuffling’ that is not seen in the outgroup lineages.

### The demographic history of *Brenthis* butterflies

To estimate the timing of rearrangements as well the effective size of the populations in which they fixed, we inferred a demographic history for the three *Brenthis* species using allele frequencies in resequenced genomes (see Methods). The best fitting demographic model estimates the *B. daphne* and *B. ino* split at 2.2 MYA and the split with *B. hecate* at 3.3 MYA (Figure 2). These speciation times allow for an estimation of the rearrangement substitution rate per genome and generation: 3.4 *×* 10*^−^*^6^, i.e. one rearrangement every *≈* 300k generations. However, assuming that rearrangements are Poisson distributed, the count of rearrangements in the population ancestral to *B. daphne* and *B. ino* (14 chromosome fusions) is significantly greater than expected given the length of the corresponding branch in the inferred species divergence history (*p <* 1 *×* 10*^−^*^4^). In other words, we can reject a model assuming a single uniform rate of rearrangement substitutions. The *N_e_*estimates of populations vary (Figure 2). The population in which the most rearrangements fixed (the ancestor of *B. daphne* and *B. ino*) has a relatively small *N_e_* (359k individuals). By contrast, the population in which the fewest rearrangements fixed (*B. ino*) has a much larger *N_e_* (1153k individuals). While this may hint at a negative relationship between *N_e_* and rearrangement rate, we cannot meaningfully test this from such a small species tree. However, these *N_e_* estimates, which are typical of insects, suggest that rearrangements do not require extremely small long-term *N_e_* to become fixed.

**Figure 2:**
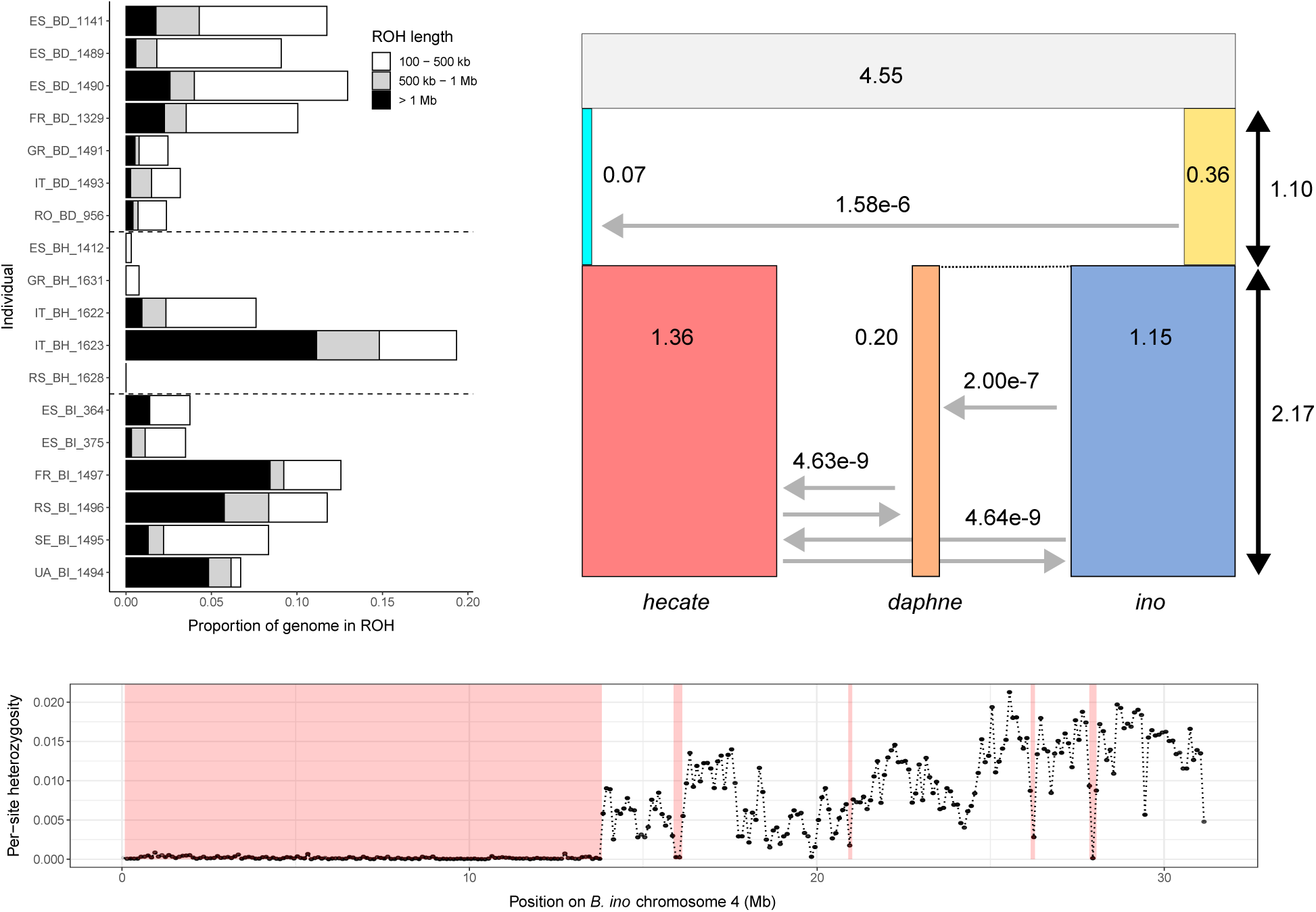
A demographic history of *Brenthis* butterflies and evidence for inbreeding. Top left: The fraction of the genome covered by runs of homozygosity (ROH) in each *Brenthis* individual. Dashed horizontal lines divide individuals by species, with *B. daphne* individuals on top, followed by *B. hecate*, and then *B. ino*. Top right: A demographic history for the three species inferred from the three dimensional site frequency spectrum (3D-SFS). Each rectangle represents a population, with width proportional to effective size (*N_e_*). Grey horizontal arrows represent asymmetrical migration between populations, whereas black vertical arrows represent the time between speciation events. Time (generations) and *N_e_* (individuals) are given in units of 10^6^. Migration arrows are labelled with their effective migration rates (*m_e_*) which are constant between speciation events. Bottom: Per-site heterozygosity for individual FR BI 1497 plotted in 100 kb windows across *B. ino* chromosome 4. Red shading shows regions of the chromosome that were identified as ROH.

We have estimated long-term *N_e_* from the SFS by modelling each species as a single panmictic population. This analysis ignores intraspecific population structure and therefore the possibility that coalescence may happen quickly within local populations but more slowly across the entire species range. With this in mind, we searched for runs of homozygosity (ROH) within individual genomes. Large ROH (*>*= 1 Mb), generated through recent shared ancestry, should be rare in well-mixed populations with *N_e_* on the order of 10^5^ individuals. Surprisingly, we found that the majority of individuals across all three species (15 of 18) have at least one ROH of this size (Figure 2). Summing the length of these ROH to estimate the inbreeding coefficient *F*_roh_ reveals that there are several individuals with *F*_roh_ *≈* 0.0625 (Figure 2), consistent with being the offspring of first cousins (which have an expected *F*_roh_ = 1*/*16).

Our inference of large long-term *N_e_* yet considerable *F*_roh_ is consistent with structured populations. For example, under the simplest model of structure – a finite-island model – local populations (i.e. demes) exchange migrants at a constant symmetrical rate and the effective size of a deme (*N_e_*) largely determines the chance of very recent common ancestry. By contrast, the longer term rate of coalescence and therefore the overall levels of genetic diversity and divergence are a function of the effective migration rate (*m_e_*) and number of demes. The parameters of this island model can be estimated numerically given windowed ROH estimates (*W*_roh_), per-base heterozygosity (*H*), and divergence (*d*_xy_) between demes (see Methods). For an *W*_roh_ (1 Mb) of 0.0221, *H* of 0.0100, and *d*_xy_ between demes of 0.0119 (which are mean values observed among *B. ino* individuals), we infer a finite-island model with 256 demes, each with an *N_e_* of 3,366, and an *m_e_* of 3.9 *×* 10*^−^*^4^. Similar parameters are estimated for *B. daphne* (demes = 20, *N_e_* = 18,794, *m_e_* = 1.3 *×* 10*^−^*^4^) and *B. hecate* (demes = 130, *N_e_*= 6,503, *m_e_*= 1.4 *×* 10*^−^*^4^). While this model is of course a drastic simplification of the likely complex population structure that exists within these dispersive species, an analogous separation of timescales is a general property of structured populations (Charlesworth *et al*. 2003; Wilkins 2004; Barton *et al*. 2010). These results shows that local populations of *Brenthis* butterflies likely have *N_e_* that is at least an order of magnitude smaller than overall diversity would suggest. Although smaller local populations may promote the fixation of rearrangements through drift, it is less clear whether this would lead to fixation across the entire species range (see Discussion).

### Evidence for selective sweeps around chromosome fusions

It is possible that the rearrangements observed in this genus have become fixed through natural selection rather than drift (see Introduction). To test this, we ask whether loci surrounding chromosome fusions show evidence for selective sweeps. The sweep inference presented in Bisschop *et al*. (2021) calculates the likelihood of a hard selective sweep given mutation counts in short sequence blocks (the bSFS). While Bisschop *et al*. (2021) used unfolded mutation counts for four lineages, this requires polarisation, i.e. knowledge of ancestral states. We can only obtain this information for genic regions of the genome given the considerable divergence between *Brenthis sp.* and the nearest available outgroup, *F. adippe* (*∼* 0.09 at 4D sites). We therefore adapted the composite likelihood based sweep inference to folded mutation counts for six lineages (Figure 3).

**Figure 3:**
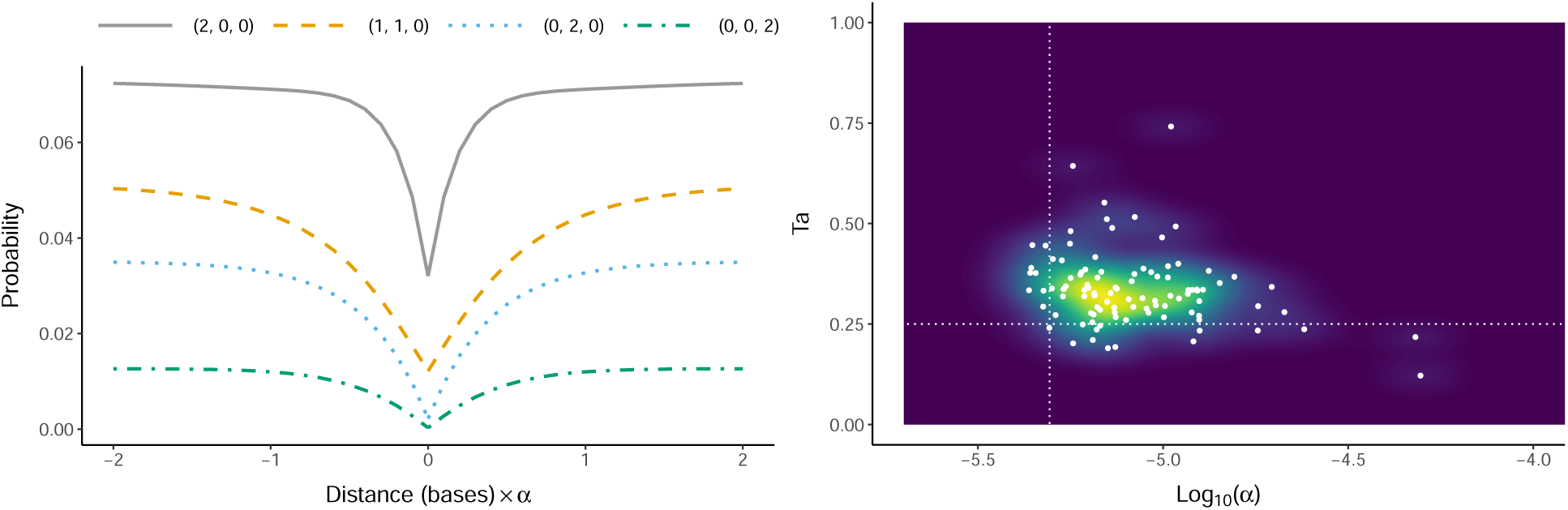
Inferring sweeps from the bSFS. Left: The probability of observing particular bSFS entries (y-axis) given the distance of a block from the sweep centre (x-axis). In this example, the sweep occurred 0.5*N_e_* generations ago (i.e. *T_a_* = 0.25) and the population mutation rate is 0.665 (e.g. *θ_per−site_* = 0.006 and block length = 113 bases). For a sample of six lineages with folded mutations and counting up to two mutations per branch type, there are 64 total bSFS entries. Each line represents one of these entries, where (*i, j, k*) denotes a block with *i* singleton mutations, *j* doubletons, and *k* tripletons. The x-axis shows that the effect of a sweep at a particular locus depends on the relative strength of the sweep (*α*) and the distance of that locus from the sweep centre. Right: Parameter estimates for 100 simulation replicates of a selective sweep. The true sweep strength (*Log*_10_(*α*), x-axis) and timing (*Ta*, y-axis) are shown with dotted vertical and horizontal lines, respectively. Each point represents parameter estimates for a single simulation replicate, and coloured contours show the density of these estimates across all replicates.

We first tested whether this implementation can accurately infer old sweeps. We simulated strong selection (*N_e_s* = 2, 500, see Methods) and estimated the statistical support for a sweep while also obtaining maximum composite likelihood estimates (MCLE) for three parameters: *θ*, *α* and *T_a_*. Here, *θ* = 4*N_e_ ∗ µ ∗ l* is the population mutation rate (where *l* is the block length), *α* = *<u>^r^</u>* ln[2*N_e_ ∗ s*] is the rate of recombination relative to the strength of the sweep, and *T_a_* is the timing of the sweep in units of 2*N_e_* generations. Across simulations, we find that the statistical support for a sweep – i.e. the increase in (log) composite likelihood (Δ ln *CL*) compared to the best fitting neutral model – is always non-zero with a median Δ ln *CL* per-block of 0.018. The population mutation rate (*θ* = 0.655) is well estimated, albeit with a small downward bias (lower quartile, median, upper quartile = 0.621, 0.635, 0.652, respectively). Similarly, the timing (*T_a_*) and strength (*α*) of the sweep are slightly overestimated and underestimated, respectively (Figure 3). These results show that, despite some bias, sweep parameters can be inferred through this method. Repeating this analysis but simulating entirely neutral evolution (see Methods) leads to inferred sweeps that are very weak or, in a minority of cases, strong but very old (Figure S2) and weakly supported: the median Δ ln *CL* across these simulations is 3.3 *×* 10*^−^*^5^ and the maximum is 0.002. Given these results, we use thresholds of *Log*_10_(*α*) *< −*4 and Δ ln *CL >* 0.002 to define plausible sweep candidates. This *α* value implies a distortion of genealogical branch lengths across at least 20 kb (Figure 3) and corresponds to *s* = 0.00017 given an *N_e_* of 1 *×* 10^6^ and a recombination rate of 2.9 *×* 10*^−^*^9^. At our chosen thresholds, we may discard some weak selective sweeps but false-positives are rare.

We next applied the same inference procedure to a total of 12 potentially recent chromosome fusions, with five, three, and four fusions sampled from *B. daphne*, *B. hecate*, and *B. ino*, respectively. Four fusions show no statistical support for a selective sweep (Table 1). The remaining eight fusions have *Log*_10_(*α*) and Δ ln *CL* values that our simulations suggest are unlikely to be observed under neutral evolution (see above). To test whether chromosome fusions have greater statistical support for selective sweeps than other regions of the genome, we fit sweep models across an entire chromosome for each species (*∼* 250 points spaced 100 kb apart per-species). We chose the same orthologous chromosome for each species - the only autosome that has not undergone any rearrangements within the genus (Figure 1). Summing the Δ ln *CL* of the 12 fusions and comparing this to points sampled from these non-rearranged chromosomes suggests that the fusions are not enriched for selective sweeps (permutation test, one-tailed p-value = 0.161). Similarly, although eight of the 12 fusions show evidence of a selective sweep, this is not a significantly different result than what can be obtained by sampling points from the non-rearranged chromosomes (permutation test, one-tailed p-value = 0.060).

The fact that we infer sweeps around some chromosome fusions, but that this is unremarkable when compared to other regions of the genome, suggests a much higher false-positive rate in the real data than in our idealised neutral simulation check. Considering points sampled across the non-rearranged chromosome, we find that 26.9% and 26.9% are classified as sweeps in *B. hecate* and *B. ino*, respectively, although the vast majority of these are old (*T_a_ >* 0.5, Figure S2). In *B. daphne* the frequency of inferred sweeps is even higher at 63.2%, and, in contrast to the other species, these sweeps are almost always estimated to be recent (*T_a_≈* 0.1, Figure S2). A plausible explanation for this is that gene flow into *B. daphne* from *B. ino* has generated genealogical histories that are better explained by a model of recent sweeps than a single panmictic population at equilibrium. Simulating data for a single population that has undergone the long-term demographic history inferred for *B. daphne* (Figure 2) and fitting a sweep model to these data, we recovered false-positive sweeps (32.0% of simulations), albeit with older inferred ages (*T_a_ ≈* 0.5, Figure S2). This suggests that gene flow and changes in *N_e_* over time likely explain at least some of the sweeps inferred around chromosome fusions.

We next consider the strength of evidence for individual sweeps around chromosome fusions. The fusion with the strongest sweep support is on chromosome 2 of the *B. daphne* genome and has a per-block Δ ln *CL* (Figure 4) that is greater than 95% of points sampled from the non-rearranged chromosome. The inferred sweep parameters (*θ* = 0.854, *Log*_10_(*α*) = -5.668, *T_a_* = 0.079) correspond to an *s* of 0.00122 and a timing of 55.7k generations ago when assuming *µ* = *r* = 2.9 *×* 10*^−^*^9^. Visualising folded i-ton counts shows a scarcity of tripleton and doubleton mutations near the fusion, as well as an overall reduction in diversity (Figure 4). In fact, there is a 264 kb region which encompasses the fusion point that does not have a single tripleton mutation (i.e. a mutation shared by three out of six lineages, Figure 4). Under a neutral model (*θ* = 0.547) the probability of a sequence block having no tripleton mutations is 0.869. By contrast, under the sweep model estimated for this region the probability of this result is 0.998, 0.996 and 0.969 for blocks sampled 1 kb, 10 kb, and 100 kb from the sweep centre, respectively.

**Figure 4:**
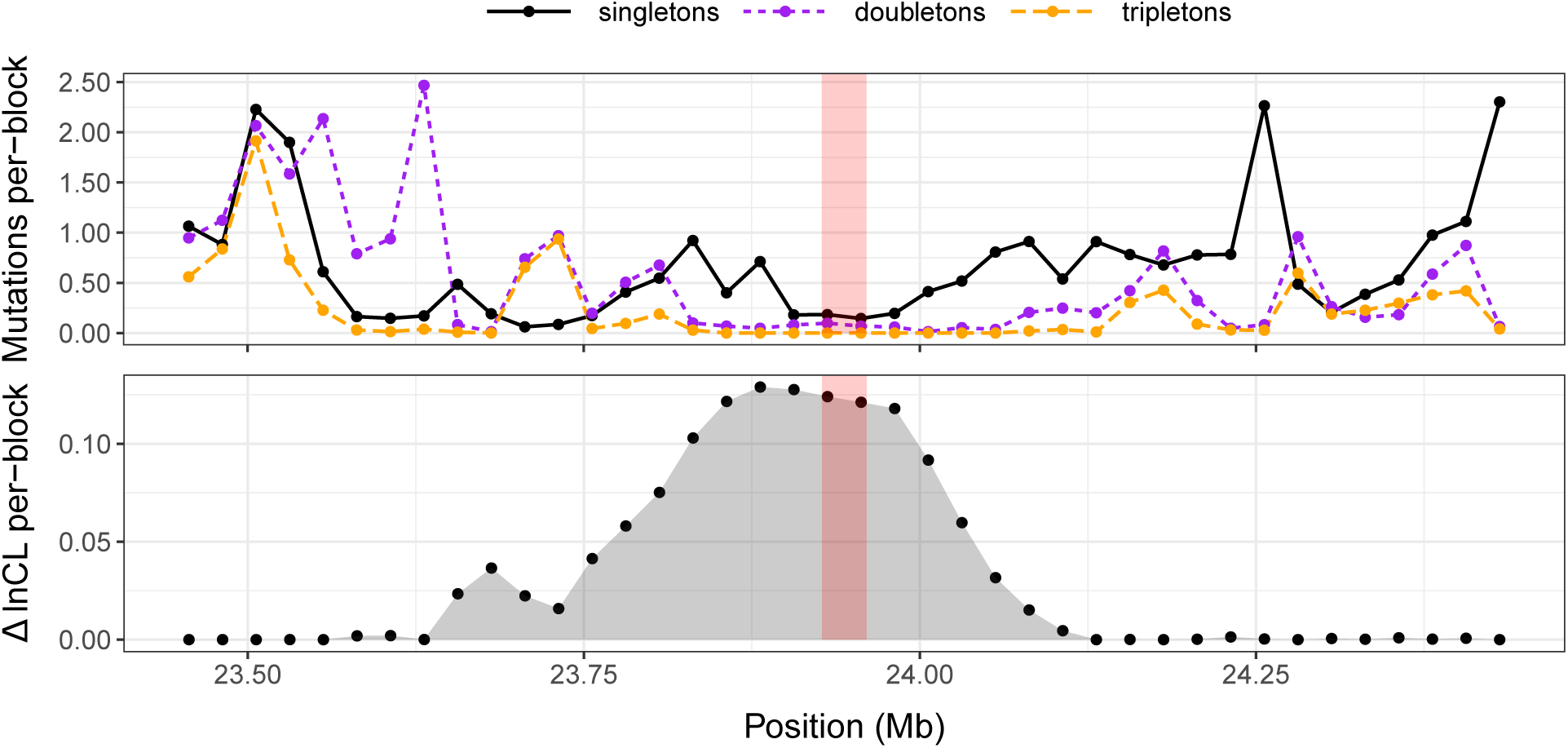
A potential selective sweep around a chromosome fusion on *B. daphne* chromosome 2. Top: The frequency (y-axis) of the three folded mutations classes across the 1 Mb of sequence that surrounds the fusion point (x-axis). Mutation frequencies are plotted in 25 kb windows, and the transparent red bar in the centre of the plot marks a 33 kb region which contains the fusion point. Bottom: The statistical support for a selective sweep (y-axis) across the same region. Support is measured as the difference in composite likelihood (Δ ln *CL*) between a selective sweep model and a neutral model. The models were fit at 40 test points at 25 kb intervals across this region, each represented by a point in the plot.

It is possible that the reduction in diversity around this chromosome fusion has been generated by processes other than a selective sweep, e.g. a lower *de novo* mutation rate or background selection. We therefore fit an alternative model to this region in which the fusion point is encompassed by a local reduction in *θ* that extends for *d* bases in either direction. We find that this model of locally reduced diversity (*θ* = 1.169, *θ_local_* = 0.391, *d* = 402 kb) fits better than the neutral model with a single *θ* parameter (per-block Δ ln *CL* = 0.110) but not as well as the sweep model (per-block Δ ln *CL* = *−*0.013). Finally, we also test whether the sweep is supported when considering all seven *B. daphne* genomes, rather than just the four originally analysed. Under this sampling, the inferred sweep is of a similar strength but older (*T_a_* = 0.26) and with reduced statistical support (per-block Δ ln *CL* = 0.057 rather than 0.123). Given the confounding effects of demography, we must interpret patterns of mutation around this chromosome fusion carefully. Nonetheless, our results do raise the possibility that this chromosome fusion has risen in frequency through a selective sweep.

## Discussion

### Patterns of chromosome evolution

Chromosome rearrangements are a fundamental part of eukaryote genome evolution, yet some groups of organisms display a much higher rate of rearrangement than others. We have focused on one such taxon, butterflies in the genus *Brenthis*, and have shown that these species have undergone a complex history of chromosome rearrangement not shared by other closely related genera (Figure 1). We find evidence for a large number of chromosome fusions as well as several fissions, with multiple rearrangements occurring between speciation events. We have assumed that chromosomes rearrange through fissions and fusions, rather than translocations. Although small segments of chromosomes appear to have translocated between *B. daphne* and *B. ino* chromosomes (Figure 1), the fact that these segments are single chromosomes in *B. hecate* suggests that these are ancestral chromosomes that have fused differentially. Overall, the pattern and tempo of rearrangement appears similar to what has been described in *Melinaea* butterflies (Nymphalidae) (Gauthier *et al*. 2022) and dissimilar from genera such as *Lysandra* (Lycaenidae) that are dominated by chromosome fissions (Wright *et al*. 2023). While it is perhaps unsurprising that the mode of rearrangement evolution differs between lineages, there do appear to be some shared features. For example, the Z sex-chromosome is one of just two chromosomes that are not rearranged between *B. hecate* and *B. daphne*, and although we have previously identified a Z-autosome fusion in one *B. ino* haplotype (Mackintosh *et al*. 2022), this rearrangement is not fixed. The Z is also the only chromosome that has not undergone extensive fissions in *Lysandra sp.*, and it is one of only two chromosomes that are not rearranged between *Melinaea marsaeus* and *M. menophilus* (Gauthier *et al*. 2022). It therefore seems likely that rearrangements involving the Z-chromosome have different fitness effects than autosomal rearrangements (Wright *et al*. 2023).

### Genetic drift and underdominance

Fissions and fusions are likely to be underdominant, i.e. deleterious when in a heterozygous state, because proper pairing and segregation of chromosomes during meiosis is often impaired (Nunes *et al*. 2011; Grize *et al*. 2019, although see Mercer *et al*. 1992; Borodin *et al*. 2019). As a result, strong genetic drift in small populations is a potential explanation for the fixation of these rearrangements (Wright 1941). We have shown that, despite high overall genetic diversity, *Brenthis* butterflies likely have much smaller local population sizes (*N_e_ ≈* 10^4^). Is it then possible that deleterious rearrangements have fixed through genetic drift in these species? Lande (1979) showed that an underdominant rearrangement which becomes established in a single deme could eventually fix across the entire population through a process of deme extinction and re-colonisation. However, this requires that the migration rate is low. Otherwise, the fixation probability within a deme diminishes (Lande 1979) and colonisation of a new deme no longer happens from a single source (Spirito *et al*. 1993). The *m_e_* values we infer under a finite-island model suggest that migration between demes is high (4*N_e_m_e_ >* 1), in which case population structure can only have a weak effect on the fixation probability of an underdominant rearrangement (Slatkin 1981). If we therefore ignore population structure and assume that *Brenthis* species typically have populations with *N_e_ ≈* 10^5^ (Figure 2), we can use the fixation rate of Lande (1979) to estimate the maximum heterozygote disadvantage of rearrangements that have fixed. Although we do not know the *de novo* rearrangement rate, we can assume that it is no higher than one rearrangement per-genome per-generation, as otherwise most individuals would be heterozygous for multiple new fissions or fusions (which we do not observe in our genome assemblies). Under this assumption, the heterozygote disadvantage *s* is *<*= 0.00014, suggesting that heterozygosity for a fission or fusion has a weak absolute fitness effect in these species. This idea is further supported by the fact that *B. daphne* and *B. ino* can produce fertile hybrids (Kitahara 2008, 2012) despite their karyotypes differing by as many as nine rearrangements (Figure 1). It is not clear how meiosis in these species is so robust to the risk of improper segregation in the presence of heterokaryotypes, although inverted meiosis is one potential explanation that has been described for other butterflies (Lukhtanov *et al*. 2018, 2020). While we cannot calculate the exact fitness effects of rearrangements in *Brenthis* butterflies, we can at least rule out the possibility that strongly underdominant rearrangements (e.g. *s >* 0.01) have fixed through genetic drift.

### The role of positive natural selection in the fixation of chromosome fusions

The scenario in which fusions are favoured by natural selection would mean that they play a role in adaptation (Yeaman 2013; Guerrero and Kirkpatrick 2014) and/or that they are driving as selfish elements (Stewart *et al*. 2019). There is currently little empirical evidence that fusions fix through positive natural selection, but this is unsurprising given that the majority of identified fusions are relatively old. For example, chromosome 2 of the human genome is the product of a fusion that happened approximately 900 kya (Poszewiecka *et al*. 2022), corresponding to *∼* 3.6*N_e_* generations in the past. Inferring the evolutionary history of such old mutations is challenging given the fact that, on average, all but two lineages in a genealogical tree coalesce within 2*N_e_* generations. We have therefore focused on species with recent chromosome fusions and a large long-term *N_e_* (Figure 2), giving us some power to detect the effects of natural selection.

We fit selective sweep models to 12 chromosome fusions and found no significant increase in statistical support for sweeps at fusion points compared to other regions of the genome. We interpret this result as evidence against a scenario where the majority of chromosome fusions fix through very strong selection, such as (holocentric) meiotic drive. It is of course possible that some chromosome fusions fix through genetic drift whereas others fix through natural selection. Only one fusion in our dataset, *B. daphne* chromosome 2 (Figure 4), has greater statistical support for a sweep than 95% of points sampled elsewhere in the genome. However, since we have considered 12 fusions, the probability that at least one meets this threshold by chance is considerable (*p* = 0.46). This fusion does, however, have greater support for a sweep than all 100 simulations performed under a *B. daphne* demography, and so sequence variation in this region cannot easily be explained by demography alone. Most, but not all, of the support for a sweep at this fusion point is due to a reduction in diversity (Figure 4). This pattern could be explained by a recombination desert in which background selection continuously erodes diversity. Although this ad-hoc explanation could be applied to almost any inferred sweep, it is at least plausible in this case as the fusion point is in the centre of a chromosome, which is where recombination is typically lowest in butterfly genomes (Shipilina *et al*. 2022; Palahí i Torres *et al*. 2022). Additionally, the fact that these fusions act as barriers to gene flow (Mackintosh *et al*. 2023) is another explanation for the reduction in diversity that we observe. Some uncertainty remains as to whether the inferred selective sweep around this chromosome fusion is a true-positive, and we therefore interpret our results as weak evidence for the idea that chromosome fusions primarily fix through positive natural selection. The patterns of mutation around this particular fusion are nonetheless unusual and so warrant further exploration. Ideally, future analyses will jointly model the effects of demography and natural selection on sequence data, which is a long-standing goal in population genomic inference (Przeworski 2002; Jensen *et al*. 2005; Lauterbur *et al*. 2022).

### Outlook

Knowledge about how genomes change over time is key for our understanding of evolution. Although fission and fusion rearrangements represent just a small fraction of the ways in which genomes can change, we know particularly little about how these drastic mutations become fixed in populations. To address this, we have analysed genome wide variation in *Brenthis* butterflies to infer past demography and natural selection in relation to chromosome rearrangements. Our main findings are that (i) drift is a stronger force than overall diversity would suggest, (ii) drift is not strong enough to fix considerably underdominant rearrangements, and (iii) there is only weak evidence that chromosome fusions fixed through positive natural selection or meiotic drive. From these results alone, we cannot yet construct a full model of how rearrangements fix in these species, but we can, however, rule out certain scenarios (see above). Clearly, other types of information not contained in genome sequence data are required for a full picture of how rearrangements fix. For example, direct estimates of heterokaryotype fitness and *de novo* rates of rearrangement are invaluable for understanding rearrangement evolution. Additionally, while we have focused on rearrangements that are likely to have fixed recently, an alternative strategy would be to identify and analyse the small subset of rearrangements that are still segregating within a species. It is more challenging to collect data on such examples but they could provide information about how rearrangements rise (and fall) in frequency over time. The population genomic analyses presented here represent a first step in understanding how fission and fusion rearrangements fix in *Brenthis* butterflies. We anticipate and look forward to similar investigations in other groups of organisms where chromosome rearrangements are common, which together will illuminate how genomes evolve across the tree of life.

## Acknowledgments

We would like to thank Sam Ebdon and Tobias Baril for kindly sharing RNA-seq and repeat annotation data, respectively. We are indebted to Vlad Dincă, Raluca Vodă and Leonardo Dapporto for contributing *B. hecate* samples and thank Dominik R. Laetsch and Alex Hayward for help with collecting the *F. adippe* sample. We would also like to thank Brian Charlesworth for comments on a previous version of this manuscript. AM is supported by an E4 PhD studentship from the Natural Environment Research Council (NE/S007407/1). KL was supported by a fellowship from the Natural Environment Research Council (NERC, NE/L011522/1). RV is supported by Grant PID2019-107078GB-I00 funded by Ministerio de Ciencia e Innovación and Agencia Estatal de Investigación (MCIN/AEI/10.13039/501100011033). SHM is supported by a Royal Society University Research Fellowship (URF/R1/180682). This work was supported by a European Research Council starting grant (ModelGenomLand 757648) to KL.

## Data accessibility and benefit sharing

All new sequence data generated in this study and the *Brenthis hecate* genome assembly are available at the European Nucleotide Archive under project accession PRJEB62818. Python scripts and *Mathematica* notebooks are available at the following Github repository: https://github.com/A-J-F-Mackintosh/Mackintosh_et_al_2023_rearrangement_fixation.

## Author contributions

AM, SHM, DS and KL designed the research. AM and DS wrote code for analysis. AM analysed genomic data. AM wrote the manuscript with input from RV, SHM, DS, and KL.

