## Supplementary material for "Do chromosome rearrangements fix by genetic drift or natural selection? A test in *Brenthis* butterflies"

### Supplementary Materials

#### Supplementary Methods

##### Gene annotation

We annotated genes in three genome assemblies: *F. adippe*, *B. daphne*, and *B. hecate* (note that *B. ino* already has a gene annotation from Mackintosh *et al.* 2022). This was done so that the SFS-based demographic modelling could be restricted to putatively neutral fourfold-degenerate (4D) sites in the *F. adippe* genome, and that exonic regions in the *Brenthis* genomes could be excluded when fitting sweep models. We masked repeats in the *B. daphne* and *B. hecate* genomes using Red (Girgis 2015) with default parameters. A repeat-masked version of the *F. adippe* assembly was kindly supplied by Tobias Baril (personal communication), having been repeat annotated with EarlGrey v1.2 (Baril *et al.* 2021, 2022).

RNA-seq data was generated for an *F. adippe* and *B. hecate* individual and kindly shared with us by Sam Ebdon (Table S1). RNA extractions, library preparations, and sequencing were performed alongside datasets generated for Ebdon *et al.* (2021). We also accessed the RNA-seq dataset for *B. daphne* from Ebdon *et al.* (2021). We next mapped species-specific RNA-seq reads to the assemblies with HISAT2 v2.1.0 (Kim *et al.* 2019). The repeat-masked assemblies and RNA-seq alignments were used as input for gene annotation with braker2.1.5 (Stanke *et al.* 2006, 2008; Li *et al.* 2009; Barnett *et al.* 2011; Lomsadze *et al.* 2014; Buchfink *et al.* 2015; Hoff *et al.* 2015, 2019). We used GenomeTools v1.6.1 (Gremme *et al.* 2013) to format gff3 and bed files for each annotation. Finally, 4D sites in the *F. adippe* genome assembly were identified with partition_cds.py (see Data Availability).

##### Fitting demographic models

We fit demographic models to the unfolded 3D_SFS using fastsimcoal2 (version fsc27093). We focused on demographic models with three current populations (*B. daphne*, *B. hecate*, *B. ino*), three ancestral populations, two speciation times with instantaneous changes in $N_e$, and eight distinct unidirectional migration rates. Because it is difficult to accurately infer this many free parameters (16 total), we took a two-step approach to identify a simple model which explains the data well and can provide accurate estimates.

We first explore the model space by allowing all parameters to vary. We then use these estimates to simplify the model and then re-run the optimization procedure. We ran 50 maximum likelihood optimisation replicates with fastsimcoal, using the options: `-n 2000000 -L 40 -w 0.001`. Across replicates, migration from *B. daphne* to *B. ino* (forwards in time) was consistently low ($\sim 2 \times 10^{-9}$). We therefore chose to set this $m_e$ parameter to zero for future model fitting. By contrast, migration in the opposite direction was much higher ($\sim 2 \times 10^{-7}$) and so this $m_e$ parameter was retained. Migration between *B. daphne* and *B. hecate*, as well as *B. ino* and *B. hecate*, were approximately symmetrical across replicates. We therefore chose to sync $m_e$ parameters in each case. Migration rates between ancestral populations were more variable.

Given these results, we considered three possible models with four $m_e$ parameters, each differing in the restriction on migration between the two ancestral populations. Model_1 has migration from the population ancestral to *B. daphne* and *B. ino* to the population ancestral to *B. hecate* forwards in time. Model_2 has migration in the opposite direction, and Model_3 has migration in both directions at the same rate. We performed maximum likelihood optimisation for each model, as described above. We found that Model_1 had the greatest composite likelihood ($-824, 681.485$), followed by Model_3 ($-825, 060.649$), then Model_2 ($-825, 090.101$). Furthermore, the composite likelihood for Model_1 is greater than the maximum observed when fitting the full model with all eight migration parameters ($-824, 695.088$). We therefore present results for Model_1 in the Main Text.

#### Fitting sweep models

We used the inference method of Bisschop *et al.* (2021) for calculating the likelihood of a selective sweep under the star-like approximation. In this method, the composite likelihood of a selective sweep is calculated by multiplying the probabilities of observing mutation configurations in short sequence blocks. Each block contains counts of (folded) singleton, doubleton, and tripleton mutations from a sample of six lineages (i.e. bSFS entries, Figure S2). The block length is consistent within each analysis and is chosen so that there are on-average 1.5 segregating sites per block. The probabilities of bSFS entries depend on the parameters of the sweep model ($\theta$, $\alpha$ and $T_a$, see Main Text) as well as the distance of a block from the sweep centre. When fitting sweep models we consider blocks within 1 Mb of a potential sweep centre, meaning that probabilities are required for many thousands of blocks. Instead of performing these calculations repeatedly, which would be prohibitively slow, we instead generated a grid with dimensions corresponding to $\theta$, $\alpha * distance$ and $T_a$, in which, each element contains the exact probabilities of all 64 possible bSFS entries. The probability of a bSFS entry for a particular parameter combination and distance from the sweep centre is obtained through linear interpolation between points in the grid. The grid contained 15 $\theta$ points between 0.1 and 1.5, 47 $\alpha * distance$ points between 0 and 12.0, and 11 $T_a$ points between 0 and 1.0. This places a limit on the age of sweeps that can be inferred ($T_a = 1$, i.e. $2N_e$ generations ago). When a sweep is weak, many blocks will be $\alpha * distance > 12$ away from the sweep centre and therefore outside of our grid. However, probabilities at such a high $\alpha * distance$ are effectively the same as under a neutral model, and we approximate the probability as such.

For a given point in the genome and the blockwise data in the surrounding 1 Mb, we optimised the parameters of the sweep model using the Nelder Mead algorithm in *Mathematica*. We repeated the optimisation three times with different random seeds and retained the parameters with the greatest likelihood. We set a minimum $Log_{10}(\alpha)$ value of -5.7. This corresponds to a very strong sweep where $\alpha *$ 500 kb = 1. Sweeps with smaller $\alpha$ values than this would be unlikely to show a spatial pattern across 1 Mb and so cannot be identified reliably.

We fit two other models to the same data. The first is a neutral model with a single parameter, $\theta$. The second is a model with a central region $2 * d$ bases in size where $\theta$ is reduced relative to a background value. Unlike the sweep model, these models do not include any distortion in genealogical branch lengths. Code for fitting all three of these models to bSFS data can be found in the *Mathematica* notebook titled brenthis_sweeps_chromosome_scan.nb (see Data Availability).

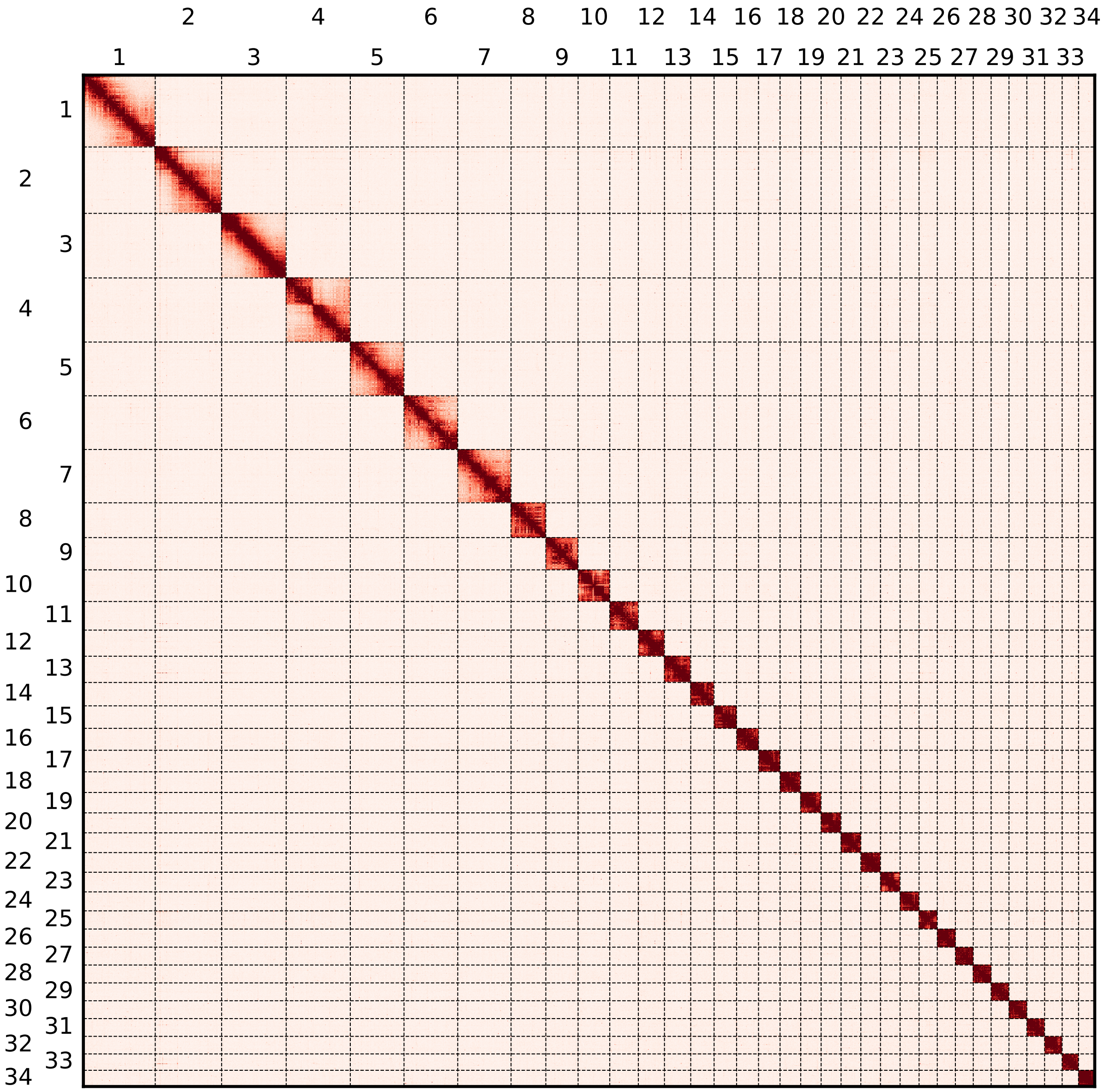


Figure S1: A HiC contact heatmap showing the 34 *Brenthis hecate* chromosomes.

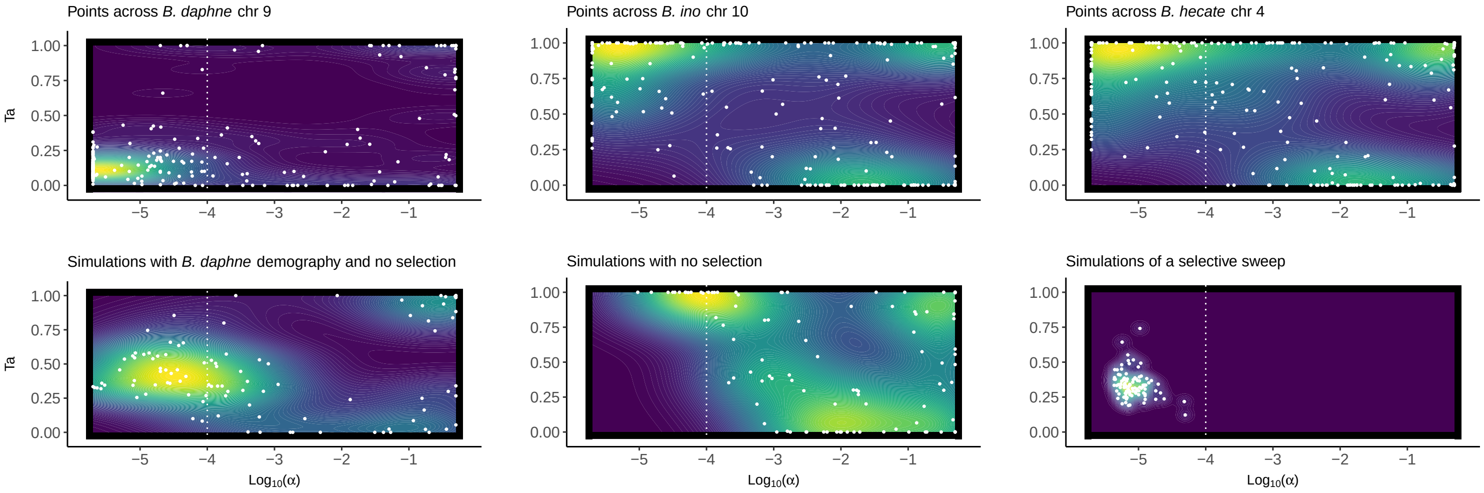


Figure S2: Parameters of inferred selective sweeps. Plots show the estimated strength of sweeps ($Log_{10}(\alpha)$, x-axis) and their estimated timing ($T_a$, y-axis). Within a plot, each white point represents parameter estimates for a test site in the genome or a single simulation, whereas coloured contours show the density of these estimates across multiple points / simulations. The top plots show inferred sweep parameters for points sampled across the same (orthologous) chromosome, for *B. daphne*, *B. ino*, and *B. hecate*. The bottom plots show inferred sweep parameters for simulations. Each plot has a vertical dashed line at $Log_{10}(\alpha) =$ -4, as points to the left of this can be considered putative selective sweeps (see Main Text).

Table S1: Sampling locations and other metadata for the individuals used to generate new sequence data in this study.

| **Sample** | **Preservation** | **Date** | **Species** | **Sex** | **Locality** | **Region** | **Country** | **Lat** | **Long** | **Collector** | **Data** |
|---|---|---|---|---|---|---|---|---|---|---|---|
| ES_BH_1411 | Liquid nitrogen | 6/6/2019 | Brenthis hecate | Male | Segura de la Sierra, Jaén | Andalucia | Spain | 38.263 | -2.615 | RV | RNA-seq, HiC |
| ES_BH_1412 | Liquid nitrogen | 10/6/2019 | Brenthis hecate | Male | Ablanque | Castille-La Mancha | Spain | 40.927 | -2.189 | RV | Pacbio, WGS |
| IT_BH_1622 | Ethanol | 14/7/2013 | Brenthis hecate | Female | Borgo Olivi | Treviso | Italy | 46.024 | 12.280 | L. Dapporto, R. Vodă | WGS |
| IT_BH_1623 | Ethanol | 22/7/2013 | Brenthis hecate | Male | Sasso Tetto | Macerata | Italy | 43.007 | 13.232 | L. Dapporto | WGS |
| RS_BH_1628 | Ethanol | 27/6/2014 | Brenthis hecate | Male | Divcibare, Mt. Maljen | NA | Serbia | 44.122 | 20.015 | R. Vodă, V. Dincă | WGS |
| GR_BH_1631 | Ethanol | 3/7/2014 | Brenthis hecate | Female | Granitis | East Macedonia and Thrace | Greece | 41.308 | 23.905 | R. Vodă, V. Dincă | WGS |
| RO_FA_934 | Liquid nitrogen | 17/7/2018 | Fabriciana adippe | Male | Pin1000m, Lupsa, Apuseni Mt. | Alba | Romania | 46.416 | 23.192 | KL, RV, Alex Hayward, Dominik R. Laetsch | RNA-seq |
